# The efficacy of commercial decontamination agents differs between standardised test settings and real-world usage for a variety of bacterial species

**DOI:** 10.1101/2022.02.08.479643

**Authors:** Benedict Uy, Hannah Read, Shara van de Pas, Rebecca Marnane, Francesca Casu, Simon Swift, Siouxsie Wiles

## Abstract

Decontamination of surfaces and items plays an important role in reducing the spread of infectious microorganisms in many settings including hospitals and research institutes. Regardless of the location, appropriate decontamination procedures are required for maintaining biosafety and biosecurity. For example, effective decontamination of microbial cultures is essential to ensure proper biocontainment and safety within microbiological laboratories. To this end, many commercial decontamination agents are available which have been tested to a prescribed standard to substantiate their efficacy. However, these standardised tests are unlikely to accurately reflect all conditions encountered in medical and biomedical research. Despite this, laboratory workers and other users of decontamination agents may assume that all decontamination agents will work in all situations. We tested commonly used commercial decontamination agents against a range of bacterial species to determine their efficacy under real-world laboratory conditions. As each decontamination agent has a different recommended dilution for use, to compare their efficacy we calculated their ‘effective ratio’ which reflects the difference between the manufacturer-recommended dilution and the dilution needed to achieve decontamination under real-world laboratory conditions. Effective ratios above 1 indicate that the agent was active at a dilution more dilute than recommended whereas effective ratios lower than 1 indicate that the agent required a higher concentration than recommended. Our results show that the quaternary ammonium agents TriGene Advance and Chemgene HLD4L were the most active out of the agents tested, with biocidal activity measured at up to 64 times the recommended dilution. In contrast, hypochlorite (bleach) and Prevail™ (stabilised hydrogen peroxide) had the lowest effective ratios amongst the tested agents. In conclusion, our data suggest that not all decontamination agents will work at the recommended dilutions under real-world laboratory conditions. We recommend that the protocols for the use of decontamination agents are verified under the specific conditions required to ensure they are fit for purpose.

## Introduction

Decontamination of surfaces and items plays an important role in reducing the spread of infectious microorganisms in many settings including hospitals and research institutes. Within microbiological laboratories, the effective decontamination of microbial cultures is essential to ensure that users are not exposed to the microorganisms being manipulated within the facility and that the microorganisms are not inadvertently released into the environment. This applies to laboratories involved in biomedical research as well as diagnostic laboratories and other testing facilities. Although each type of facility will have different samples and decontamination applications, all will have guidelines to prevent and reduce the occurrence of containment breaches. Proper decontamination procedures are a part of these guidelines.

There are many options for decontaminating contaminated materials. Autoclaving is one such method, however it is unsuitable for heat-sensitive items, and depending on the size of the autoclave, for large items. It is also inappropriate for decontaminating some fixed items, such as bench top surfaces. Where autoclaving is not an option, chemical decontamination agents can be used. Six main classes of chemical decontamination agents are routinely used: iodophors, quaternary ammonium compounds (QAC), peroxides, phenols, chlorine, and aldehydes (Chapman, 2003). Each class has a different mechanism of action and their own set of advantages and disadvantages (Geraghty et al., 2014). The type of agent that is used can also depend on the application; for example, aldehydes are generally used for the decontamination of medical devices, while iodophors are generally used as antiseptics (Bell et al., 1989).

Decontamination agents of the different classes are commercially available, and companies selling these agents verify the bactericidal claims through independent standardised testing by certified laboratories. The specific standards for these tests include a pre-determined set of conditions used to test their efficacy against specific microorganisms. These standards can be for a continent such as the European Standards (EN) or a specific country such as the American Society for Testing and Material (ATSM), and the Environmental Protection Agency (EPA) standards for the United States, or the Canadian standards from the Standards Council of Canada. There are also international standards such as those from the Association of Official Analytical Chemists (AOAC International) and the Clinical & Laboratory Standards Institute (CLSI). For each standard, there are separate bactericidal, sporicidal, mycobactericidal, and fungicidal tests. For example, European Standard EN 1276 is the standard for evaluating the bactericidal activity of chemical disinfectants and antiseptics used in food, industrial, domestic and institutional areas, whereas EN 1650 is the fungal equivalent. The standards also differ in their purpose; EN 1656 is tailored for determining bactericidal activities in veterinary areas, while EN 13623 investigates bactericidal activity in aqueous systems. Similarly, EN 1276 is appropriate for testing materials in suspension whereas EN 13697 is appropriate for non-porous surfaces.

While the name of each standard offers an indication of the testing aims, the detailed protocols are often hidden behind a paywall making it difficult to ascertain how each test is performed. Despite this, decontamination agent users may assume that the conditions used for each standard by each company would be the same and thus comparable. However, this is not the case. For example, according to their websites, the companies that manufacture TriGene Advance and Chemgene HLD4L report that testing activity against *Mycobacterium terrae* was performed using different dilutions, holding times, and endpoints despite both using the EN 14348 standard. This is concerning as hold time and dilution are known to be important in determining the efficacy of decontamination (Rutala & Weber, 1999; Russell & McDonnell, 2000). Although not stated, it could be assumed that the inocula for both tests were similar as microbial load is known to influence decontamination efficacy (Rutala & Weber, 1999; Rutala & Weber, 2015). Also concerning is that the Chemgene product label currently recommends a lower dilution and shorter hold time for decontaminating mycobacterial-contaminated items compared to those they tested for determining efficacy.

One other important consideration is whether the endpoints used within the standards to determine antimicrobial activity is relevant for microbiological laboratories. For example, the EN 1276 bactericidal tests use a >5-log reduction to determine activity. However, when working with pure cultures of bacteria in a research laboratory, concentrations of 10^9^ colony forming units (CFU) per mL are often reached with total volumes ranging from less than 5 mL to several litres. Under optimum conditions, many fast-growing bacterial species can reach these concentrations within 24 hours. This means that to be effective in practice, decontamination agents must be able to kill a larger number of cells and achieve more than the 5-log reduction set in the EN 1276 standard.

In this study, we investigated the efficacy of commonly used and commercially available decontamination agents against a variety of pathogens, including Gram-positive, Gram-negative, and Mycobacterial spp. Importantly, we designed our tests to mimic the decontamination of contaminated liquid laboratory wastes generated in standard microbiological research laboratories. In addition, we investigated the efficacy of these decontamination agents against antibiotic-resistant strains of *Escherichia coli* which we hypothesised could be more resistant to killing by the decontamination agents due to cross resistance (Tattawasart et al., 1999; Wesgate, Grasha & Maillard, 2016). Our results show that not all decontamination agents will work at the recommended dilutions under real-world laboratory conditions.

## Materials & Methods

### Bacterial strains

The bacterial strains used in this study are shown in Table 1. Bacteria were revived from frozen stocks stored at -80 °C prior to use to prevent adaptation over multiple laboratory subcultures. Bacteria were grown to stationary phase using the appropriate liquid media and temperature (Table 1). All growth media and supplements were purchased from Fort Richard (New Zealand). Liquid cultures of mycobacterial species were grown in Middlebrook 7H9 broth supplemented with 10% Middlebrook ADC enrichment media, 0.4% glycerol (Sigma-Aldrich, New Zealand), and 0.05% Tween 20 (Sigma-Aldrich, New Zealand). *Mycobacterium smegmatis* and *M. abscessus* were grown at 37 °C for 42-48 h while *M. marinum* was grown at 28 °C for 10-14 days. *M. tuberculosis* was grown for 4-6 weeks at 37 °C in a Biosafety Level 3 (BSL3) laboratory. All other cultures were grown for 18-24 h. Stationary phase Mycobacterial cultures were adjusted to an optical density at 600 nm (OD_600_) of 1 in 7H9 broth supplemented with 10% ADC enrichment media. All other bacteria were adjusted to an OD_600_ of 1 in BD BBL^MT^ cation-adjusted Mueller-Hinton II Broth (MHB). Adjusted cultures were serially diluted and plated on solid media to determine the initial inocula, with concentrations ranging from 2 ×10^7^ to 5 ×10^9^CFU/mL, depending on the organism. For Mycobacterial cultures Middlebrook 7H10 agar supplemented with 10% Middlebrook OADC enrichment media was used. All other bacteria were grown on Mueller-Hinton II Agar.

**Table 1.** Bacterial Strains used in this study.

| Species | Strain | Description | Growth Medium | Temperature (°C) | Ref |
| --- | --- | --- | --- | --- | --- |
| *Acinetobacter baumannii* | BAA-1790 | Multidrug-resistant; isolated from sputum | Tryptic Soy | 37 | Korotetskiy *et al* (2020) |
| *Bacillus spizizenii* | ATCC 6633 | Formerly *B. subtilis*; quality control strain | Brain Heart Infusion | 30 | ATCC |
| *Escherichia coli* | ATCC 25922 | Quality control strain | Tryptic Soy | 37 | Minogue *et al* (2014) |
|  | BAA-2340 | Carbapenem-resistant KPC reference strain | Nutrient | 37 | Trepanier *et al* (2017) |
|  | BAA-2452 | Carbapenem-resistant NDM-1 reference strain | Tryptic Soy | 37 | Trepanier *et al* (2017) |
|  | Top 10 | Antibiotic-susceptible molecular cloning strain | Luria-Bertani | 37 | ThermoFisher Scientific |
| *Klebsiella oxytoca* | ATCC 49131 | Quality control strain | Tryptic Soy | 37 | ATCC |
| *Mycobacterium abscessus* | NZRM 4048 | Clinical isolate | 7H9 | 37 | Freeman *et al* (2007) |
| *Mycobacterium marinum* | ATCC 927 | Type strain; isolate from fish | 7H9 | 30 | Aronson (1926) |
| *Mycobacterium smegmatis* | ATCC 700084 | Sensitive to kanamycin; mc<sup>2</sup>155 | 7H9 | 37 | Snapper *et al* (1990) |
| *Mycobacterium tuberculosis* | ATCC 25618 | Virulent; H37Rv | 7H9 | 37 | Oatway & Steenken (1936) |
| *Pseudomonas aeruginosa* | ATCC 27853 | Quality control strain | Tryptic Soy | 37 | Cao *et al* (2017) |
| *Staphylococcus aureus* | ATCC 6538 | Quality control strain | Tryptic Soy | 37 | Makarova *et al* (2017) |

### Decontamination agent preparation

Commercial decontamination agents were purchased from their respective suppliers (Table 2) and were tested at a wide range of concentrations which included their recommended dilution. The agents were diluted in 7H9 broth supplemented with 10% ADC for the mycobacterial strains, and in MHB for all other bacteria.

**Table 2.** Decontamination agents used in this study.

| **Decontamination agent** | **Manufacturer** | **Active component** | **Recommended dilution** |
| --- | --- | --- | --- |
| Bleach<sup>1</sup> | Brighton Professional | Sodium hypochlorite | 1:10 |
| HLD4L | Chemgene | Surfactant with quaternary ammonium compounds | 1:100 |
| Prevail<sup>TM</sup> | Virox | Hydrogen peroxide | 1:40 |
| TriGene Advance | Trisiel | Poly biguanide with quaternary ammonium compounds | 1:100 |
| Virkon | Dupont | Oxone, benzenesulfonate and sulfamic acid mixture | 1:100 (as a 1% w/v solution) |

<sup>1</sup>4% solution of stabilised liquid bleach with 53g/L of hypochlorite.

### Minimum Bactericidal Dilution Factor (MBDF) Assay

Minimum Bactericidal Dilution Factors (MBDF) were determined using microdilutions in a microtiter plate. We added 200 µL of decontamination agent in duplicate to the top wells of a clear, flat-bottomed 96-well plate (Nunc, Thermo Scientific, New Zealand) and filled the remaining wells with 100 µL of either 7H9 for mycobacteria or MHB for all other species. 100 µL of each decontamination agent was then serially diluted down the 96-well plate to give doubling dilutions. If greater dilutions were needed, the agent was initially diluted in the appropriate media before adding to the top wells of the microtiter plate. Then we added 100 µL of diluted bacterial culture to each well and incubated the plates static at room temperature. At various time points over four hours, 4 × 10µL aliquots were removed from each well and plated onto the appropriate agar plates. These were incubated at the appropriate temperature/time combination depending on the bacterial strain being tested and any growth determined by eye. The MBDF was calculated as the greatest dilution of disinfectant returning no visible bacterial growth.

### Effective ratio calculation

Due to the different recommended dilutions for the decontamination agents, an Effective Ratio was calculated by dividing the MBDF by the recommended dilution (Table 2). An effective ratio of 1 indicates the agent is active at the recommended dilution. A ratio greater than 1 indicates that the decontamination agent is active at a dilution more dilute than recommended. An effective ratio lower than 1 indicates that the agent is not active at the recommended dilution.

### Statistical analysis

Statistical analysis was performed using GraphPad Prism version 8.4.3. Data was analysed using a mixed-effects model with Tukey’s multiple comparison test. Statistical significance was set at P ≤ 0.05.

## Results

We investigated the bactericidal activity of bleach, Chemgene HLD4L, Prevail™, TriGene Advance, and Virkon against a variety of Gram-positive, Gram-negative, and Mycobacterial species. The minimum dilutions we observed as necessary for bactericidal activity (given as median and range) are presented in Table 3. Due to the decontamination agents having different recommended dilutions for use (Table 2), to easily visualise the efficacy of each agent we calculated an Effective Ratio for each bacterium-decontamination agent combination (Fig 1.).

**Table 3.**
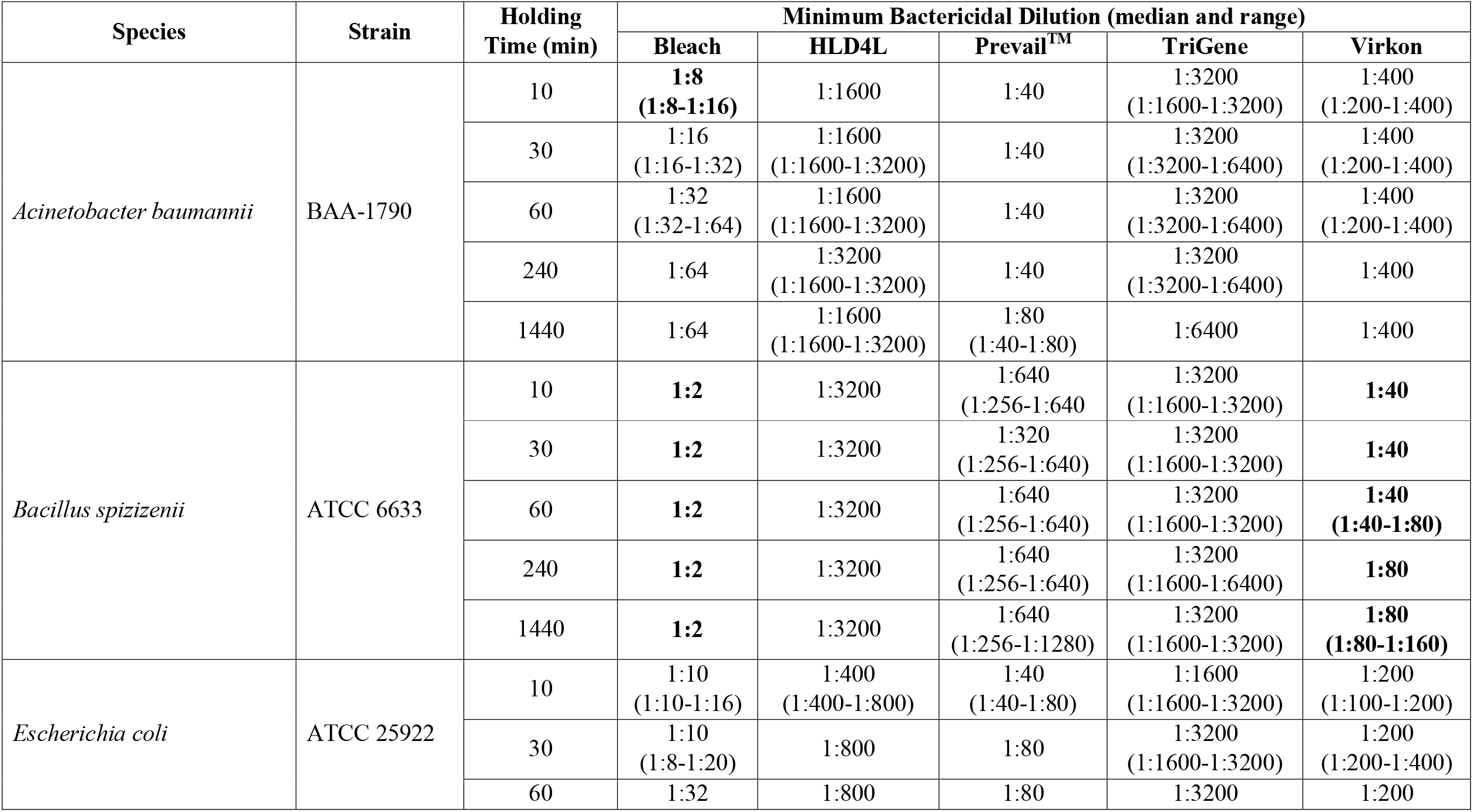
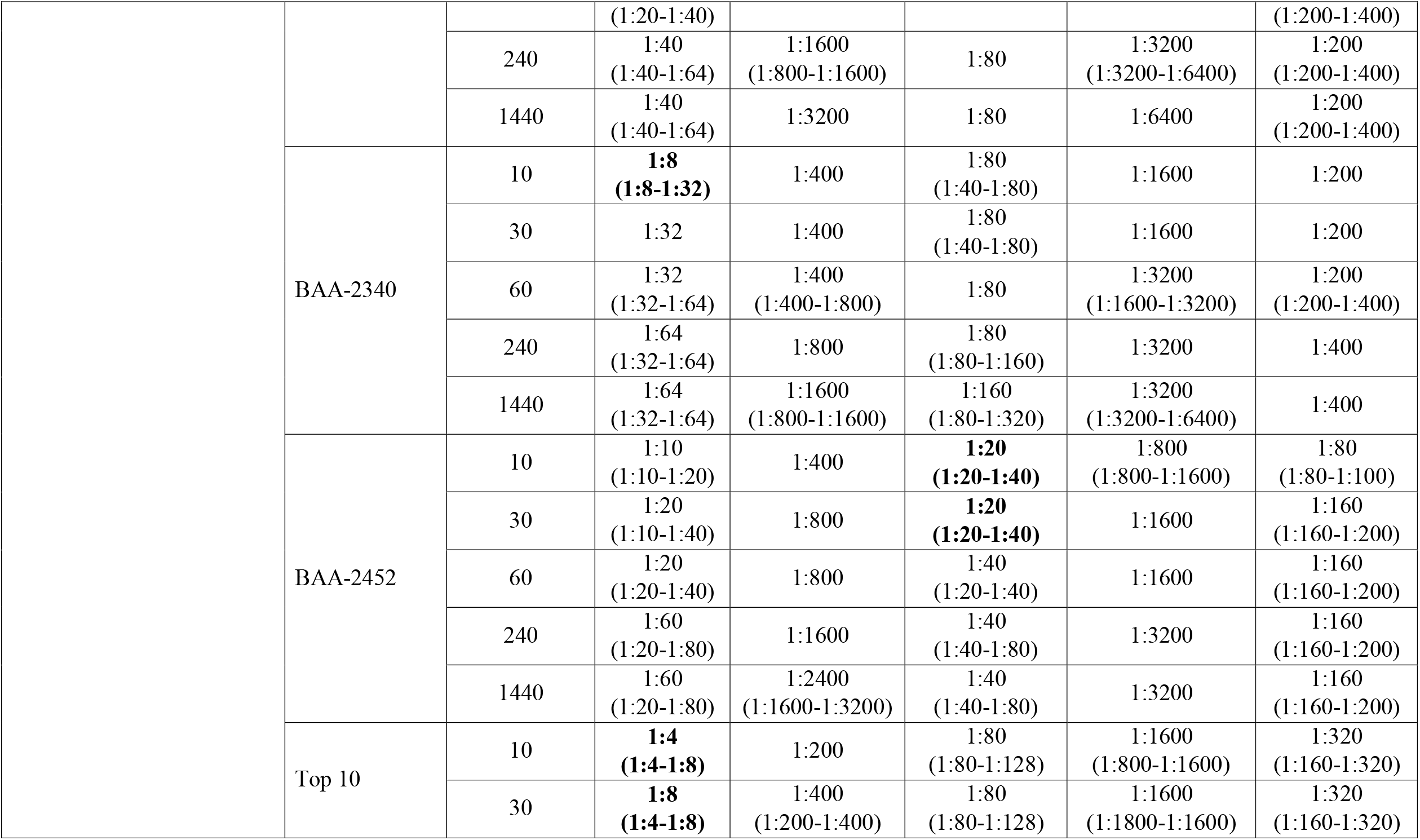
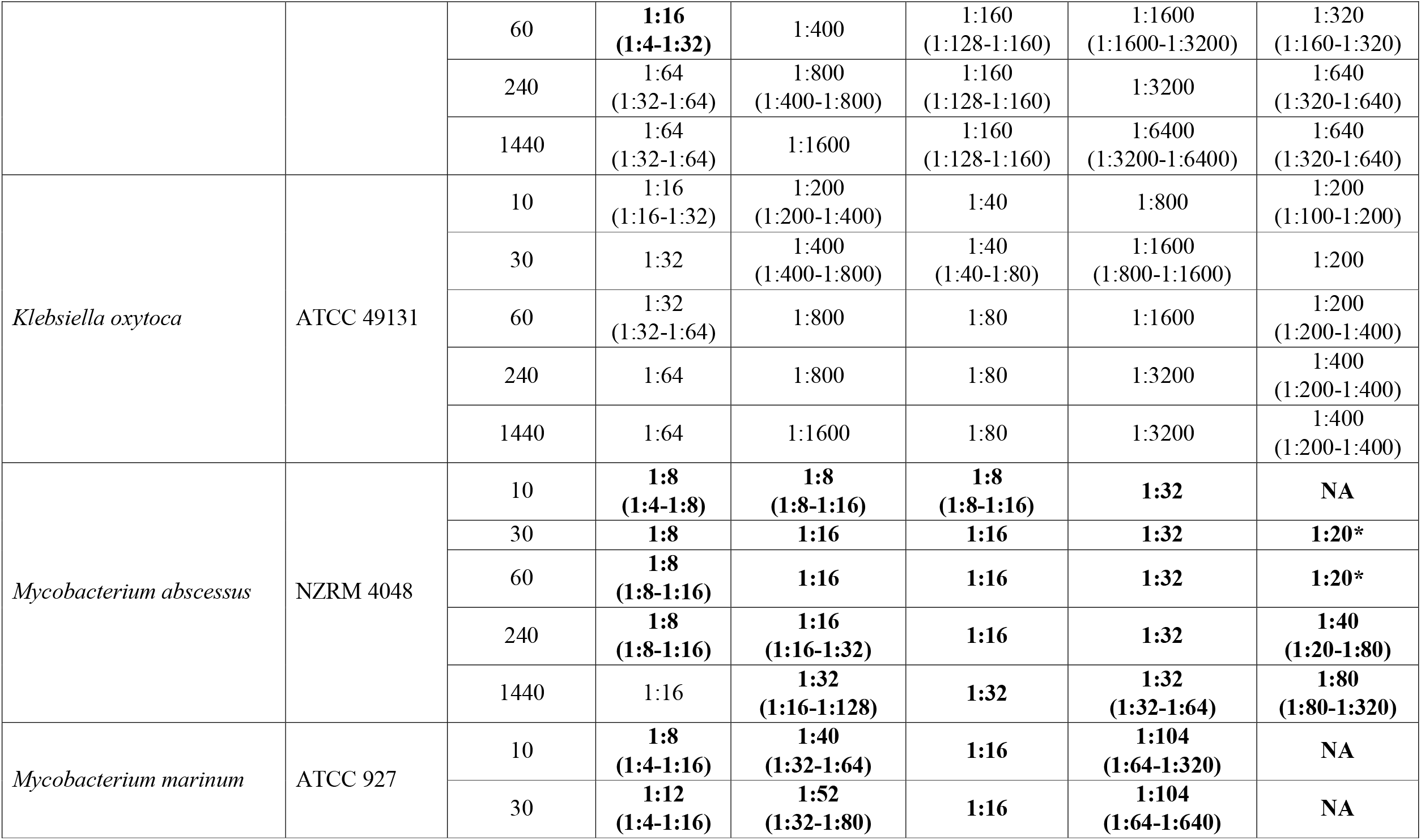
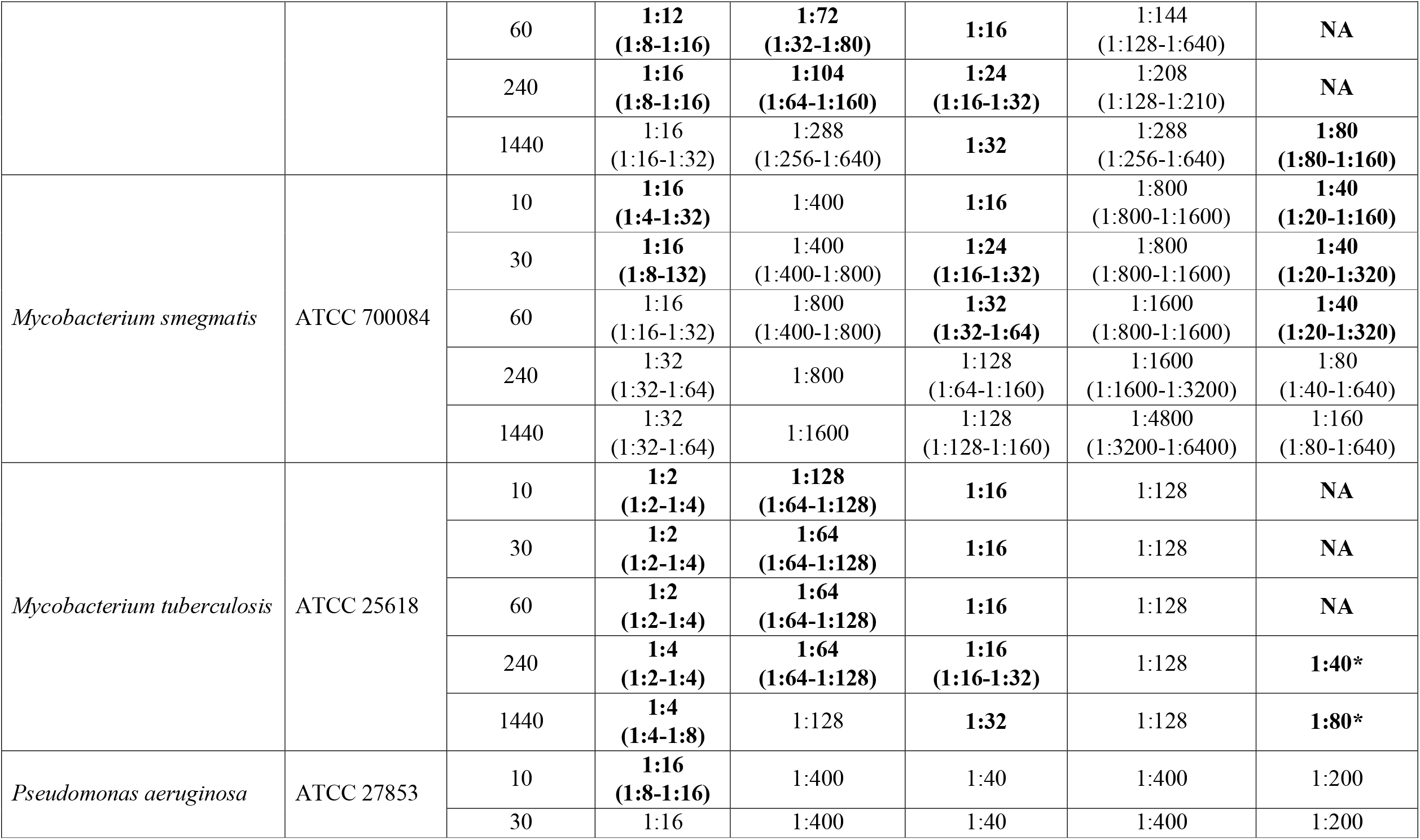
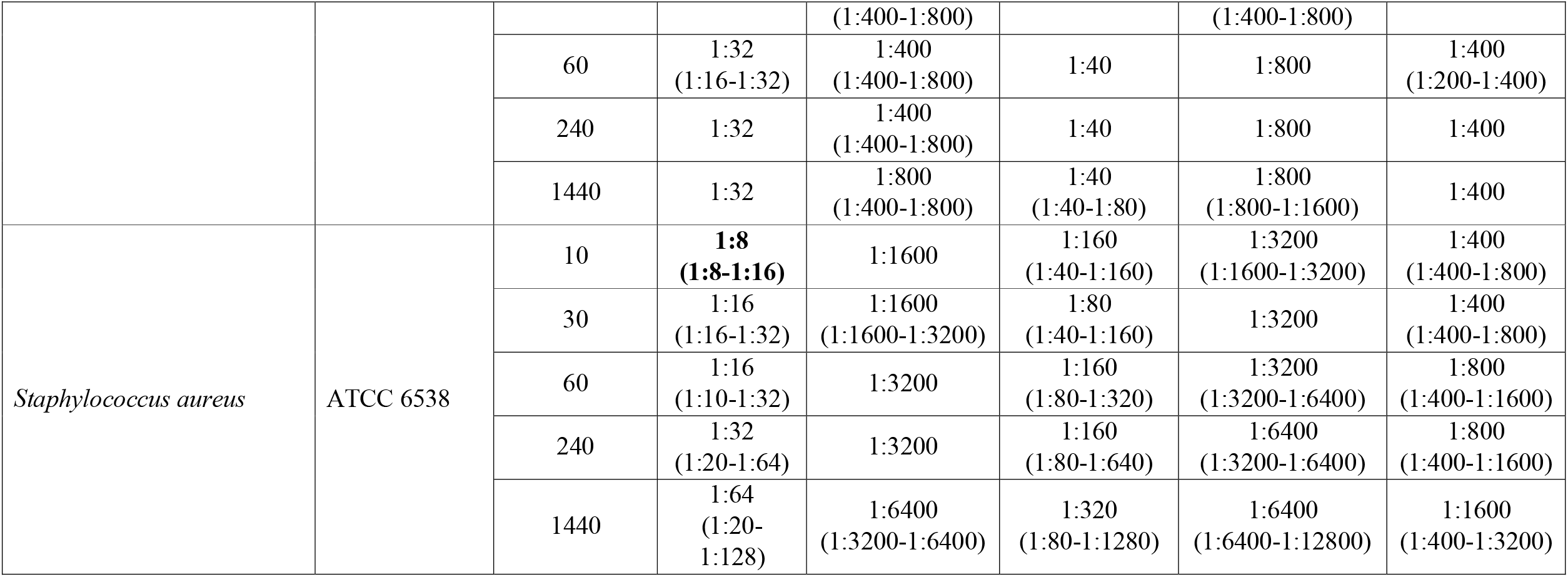
Minimum Bactericidal Dilution for various decontaminating agents at different holding times. Values shown in bold indicate a condition under which the recommended dilution would not be sufficient for decontamination.

**Figure 1.**
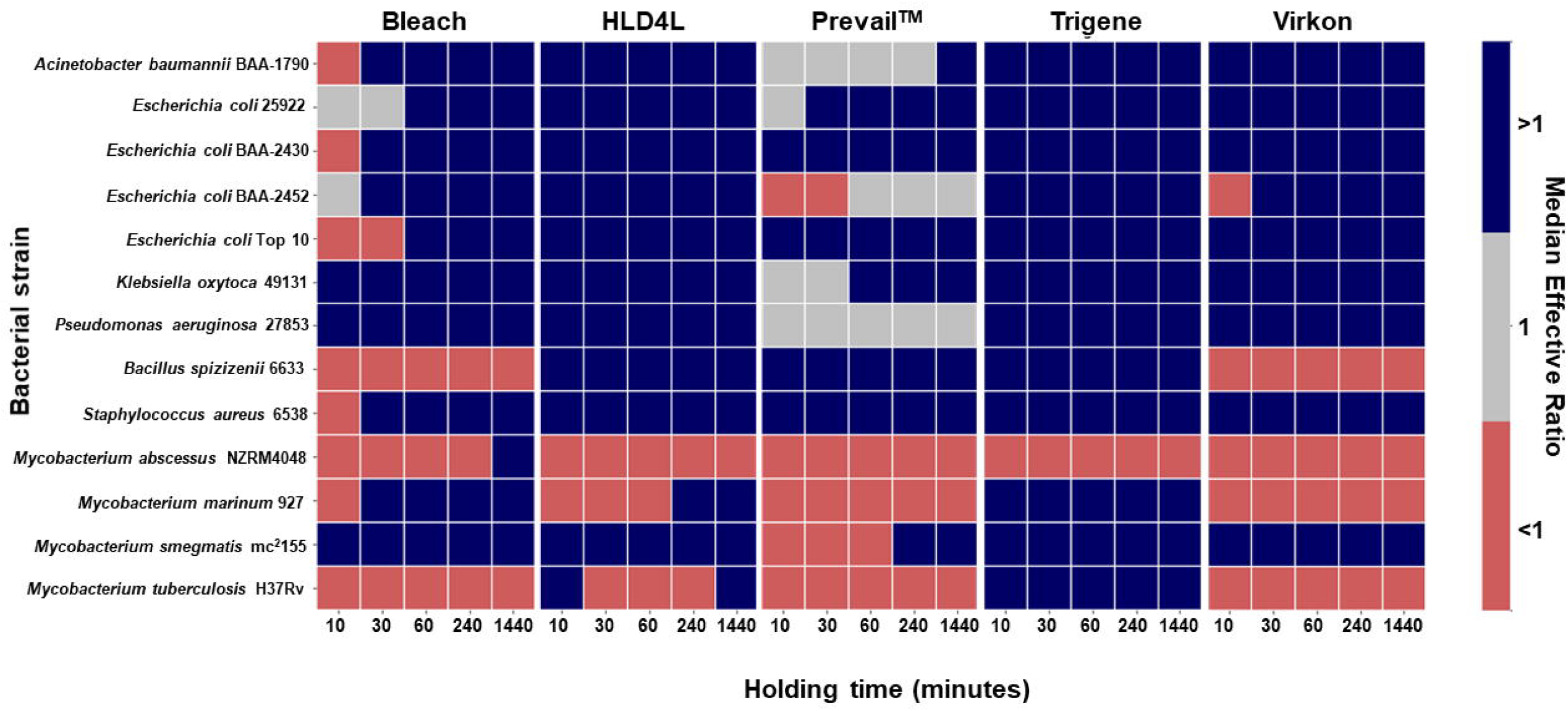
Effective ratios of various decontamination agents against a range of bacterial species at different hold times. As each decontamination agent has a different recommended dilution for use, to compare their efficacy we calculated their effective ratios by dividing the Minimum Bactericidal Dilution Factor by the recommended dilution for each agent. An Effective Ratio (ER) of 1 (grey boxes) indicates an agent active at the recommended dilution, while an ER above 1 (dark blue boxes) indicates an agent was active at a dilution more dilute than recommended and an ER below 1 (orange boxes) indicates an agent that was not active at the recommended dilution. Data (n=3-5) is presented as the median ER for each species at a given hold time. The raw data is available online from https://doi.org/10.17608/k6.auckland.19142606.

This was achieved by dividing the minimum bactericidal dilution by the recommended dilution. An Effective Ratio of 1 indicates an agent is active at the recommended dilution. A ratio greater than 1 indicates that an agent is active at a dilution more dilute than recommended while a ratio lower than 1 indicates that an agent is not active at the recommended dilution.

### Activity of decontaminating agents against Gram-negative bacteria

We used *A. baumannii, E. coli, K. oxytoca*, and *P. aeruginosa* as representative Gram-negative organisms. As can be seen from the data provided in Table 3 and Fig 1, Chemgene HLD4L, TriGene Advance, and Virkon were bactericidal against *A. baumannii, K. oxytoca*, and *P. aeruginosa* with Effective Ratios above 1 showing that these agents were active at dilutions greater than those recommended at all holding times (Fig. 1). With Effective Ratios of 1, Prevail™ was active against these species at the recommended dilution at holding times as short as 10 min (Fig. 1). While bleach was active against *K. oxytoca* at the recommended dilution for all holding times tested, in some experiments with *A. baumannii* and *P. aeruginosa*, Effective Ratios were less than 1 indicating that a lower dilution than that recommended was needed to achieve bactericidal activity using a holding time of 10 min (Fig. 1).

Given the increased prevalence of antibiotic-resistant bacteria, we investigated whether the decontamination agents would have similar activity against *E. coli* strains with different resistance profiles. ATCC 25922 is an antibiotic-sensitive strain recommended for quality control purposes while Top 10 is a commonly used molecular cloning strain that is also sensitive to antibiotics. In contrast, BAA-2340 and BAA-2452 are both carbapenem resistant. According to the ATCC website, BAA-2340 encodes a *Klebsiella pneumonia* carbapenemase (KPC) while BAA-2452 encodes a New Delhi Metalloprotease 1 (NDM-1). Overall, the decontamination agents were effective against the four *E. coli* strains with some minor differences. Bleach was effective at the recommended dilution except when used against BAA-2340 with a 10 min holding time and Top 10 with holding times of 10, 30, and 60 min (Table 3, Fig. 1). Prevail™ also required a lower dilution than recommended when used against BAA-2452 with hold times of 10, 30, and 60 min.

We also calculated Area Under Curve values from the minimum bactericidal dilutions obtained for the *E. coli* strains over the five holding times (Fig. 2). Lower AUC values reflect lower dilutions required for activity and hence increased resistance of the bacterium to the decontamination agent. We analysed this data using a mixed-effects model with Tukey’s multiple comparison test. The results indicate that Top 10 is significantly more resistant to HLD4L than ATCC 25922 (p = 0.0188), BAA-2452 is significantly more resistant to Trigene than ATCC 25922 (p = 0.0346), and that BAA-2452 is significantly more resistant to Virkon than BAA-2430 (p < 0.0001).

**Figure 2.**
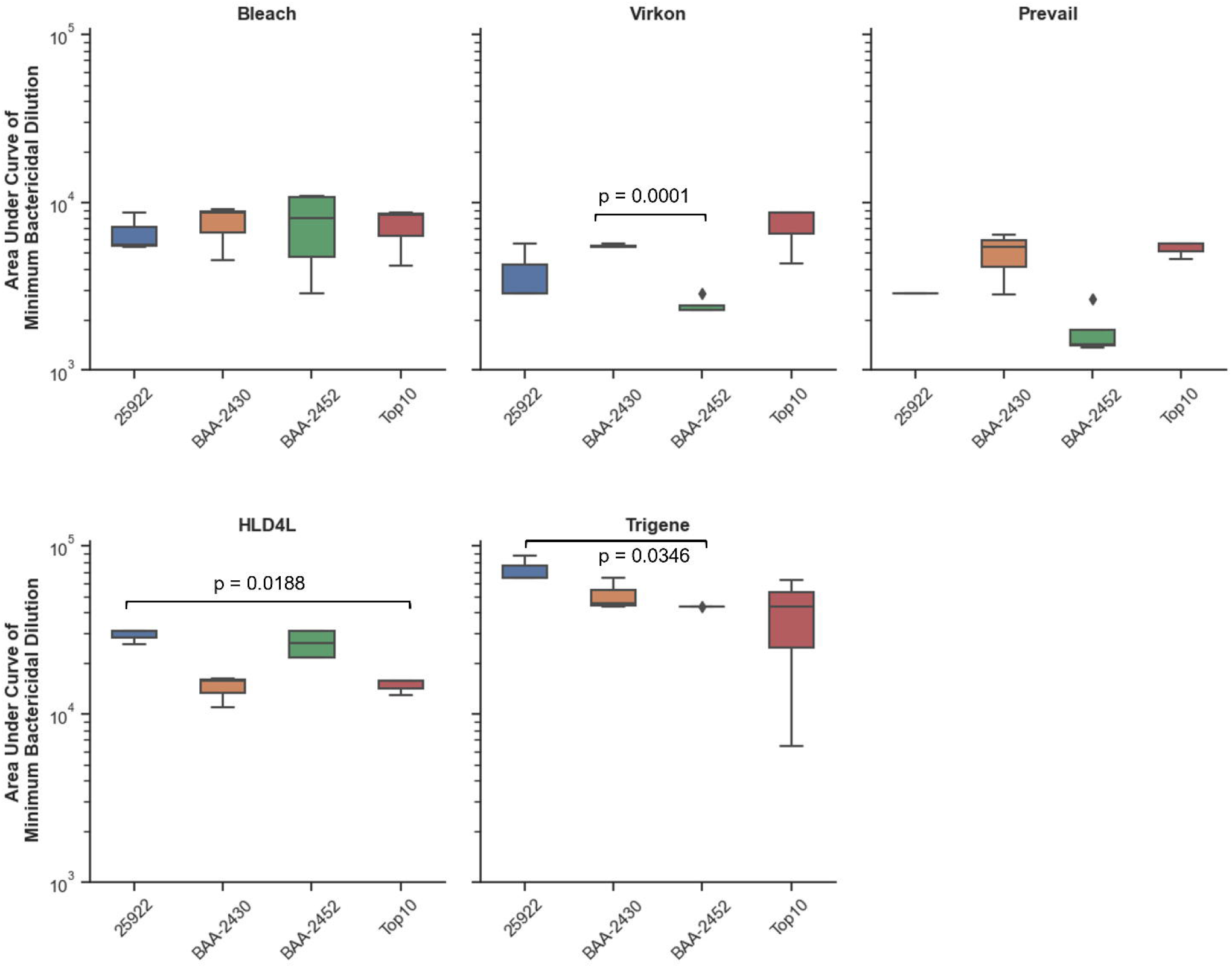
The efficacy of decontamination agents can vary between *E. coli* isolates To compare the efficacy of the decontamination agents between different *E. coli* isolates (ATCC 25922 [antibiotic-sensitive]; BAA-2340 [carbapenem-resistant, KPC^+^]; BAA-2452 [carbapenem-resistant, NDM-1^+^]; Top 10 [antibiotic-sensitive, molecular cloning strain]) we calculated Area Under Curve (AUC) values from the minimum bactericidal dilutions obtained over the five holding times. Lower AUC values reflect lower dilutions required for activity and hence increased resistance of the bacterium to the decontamination agent. Data (n=3-5) is presented as box and whisker plots with median. The data was analysed using a mixed-effects model with Tukey’s multiple comparison test. The raw data is available online from https://doi.org/10.17608/k6.auckland.19142606.

### Activity of decontaminating agents against Gram-positive bacteria

We used *B. spizizenii* (formerly *B. subtilis* subspecies *spizizenii*) and *S. aureus* as representative Gram-positive species. *B. spizizenii* is also a spore-forming bacterium. As can be seen from the data provided in Table 3 and Fig 1, Chemgene HLD4L, Prevail™, and TriGene Advance were bactericidal against *B. spizizenii* at dilutions far exceeding the recommended dilution at all holding times. In contrast, with Effective Ratios less than 1, bleach and Virkon required a lower dilution than that recommended to achieve bactericidal activity at all holding times (Fig. 1). For *S. aureus*, all decontamination agents were effective at or below the recommended dilution except for bleach with a holding time of 10 min. There was also a clear relationship between concentration and time, with all decontaminating agents being effective against S. aureus at higher dilutions as the holding time increased.

### Activity of decontaminating agents against Mycobacteria

We tested four Mycobacterial species, *M. abscessus, M. marinum, M. smegmatis*, and *M. tuberculosis*, against the panel of decontamination agents (Table 3, Fig. 1). None of the agents was active against *M. abscessus* at the recommended dilution, except for bleach at a holding time of 1440 min. The data is similar for *M. marinum*, with the addition of Trigene Advance being active at the recommended dilution for holding times of 60 min or more. *M. smegmatis* was the most sensitive of the four mycobacterial species tested, with both Chemgene HLD4L and Trigene Advance active at or higher than the recommended dilution at all holding times. The remaining decontamination agents were also active at or higher than the recommended dilution at holding times at 240 min and longer except for bleach which was also active at 60 min. For *M. tuberculosis*, Trigene Advance was the decontamination agent active at or higher than the recommended dilution at all holding times. Chemgene HLD4L was also active above the recommended dilution but only with a holding time of 1440 min.

## Discussion

It is reassuring that many of the decontamination agents tested work above the parameters assessed in the standards against a wide range of bacterial species. Even if the decontamination agent works at a higher efficacy than is recommended by the manufacturer, it is still prudent to use the recommended dilution to ensure complete sterilisation. However, for some organism and decontamination agent combinations, we obtained an effective ratio below one indicating that the recommended dilution was not active.

In many situations, the real-world usage of decontamination agents will differ from the conditions they were tested under using the standards. In the worst-case scenario, this will result in a decontamination failure. For example, Prevail™ is advertised as an anti-mycobacterial agent with activity against *M. tuberculosis* with a contact time of just 5 min. Yet our data shows that it was only active at the recommended dilution against the non-pathogenic *M. smegmatis* at holding times of 240 min or greater and was not effective against pathogenic species such as *M. tuberculosis* and *M. abscessus*. The enhanced resistance of Mycobacteria against many chemical agents is well known and has led to the development of separate standards for testing against Mycobacteria (Best et al., 1990; Griffiths, Babb & Fraise, 1998; Russell, 2001). Standards EN 14204 and EN 14348 evaluate the mycobactericidal activity of chemical disinfectants and antiseptics used in the veterinary area and in the medical area, respectively. One reason for the discrepancy between our results and the advertised activity of Prevail™ may be because *M. smegmatis* has less stringent nutrients requirements for growth so the testing conditions may not have recapitulated the standard growth conditions of other, arguably more important, species. Supplementation with catalase is common when growing many mycobacterial species including *M. tuberculosis*. As catalase mitigates against the toxic effects of hydrogen peroxide by converting it to water and oxygen, it is unsurprising this would have an impact on the activity of Prevail™ which is a hydrogen peroxide-based decontamination agent.

Our data serves as a warning that decontamination agents may not be effective against all strains of a particular species. In our study, we tested the efficacy of five decontamination agents against four strains of *E. coli*, including two antibiotic-sensitive and two antibiotic-resistant isolates. Biocides generally have non-specific targets which eventuate in cell death. As with other antimicrobials, resistance to decontamination agents can develop. Indeed, many organisms can develop natural resistance against certain compounds, such as how Pseudomonads have generally high intrinsic resistance against many agents including antibiotics (Adair, Geftic & Gelzer, 1969; Aires et al., 1999). Resistance or tolerance to a decontamination agent could have a natural genetic basis or be acquired through co-resistance with antibiotic resistance genes. Tattawasart and colleagues showed that some chlorhexidine-resistant strains of *Pseudomonas stutzeri* have cross-resistance against other biocides such as triclosan, as well as antibiotics including rifampicin and polymyxin B (Tattawasart et al., 1999). This cross-resistance between biocides and antibiotics was also identified with the use of triclosan and prolonged hydrogen peroxide treatment (Tattawasart et al., 1999; Wesgate, Grasha & Maillard, 2016).

Our data shows that the antibiotic-resistant *E. coli* strain BAA-2452 is more resistant to TriGene Advance than the antibiotic-sensitive quality control strain ATCC 25922, and more resistant to Virkon than the antibiotic-resistant strain BAA-2430. BAA-2340 and BAA-2452 are both carbapenem resistant though they encode different carbapenemases. Resistance in BAA-2452 is attributed to the acquisition of the *Klebsiella pneumoniae* Carbapenemase (KPC) encoded by *bla*_KPC_. The KPC was initially acquired through transposon Tn4401 which encoded a β-lactamase that can hydrolyse carbapenems, a class of β-lactam based antibiotics resistant against degradation by other β-lactamases (Cuzon, Naas & Nordmann, 2011). In this case, it is unlikely that a β-lactamase would be involved in resistance against a quaternary ammonium decontamination agent. Bacterial membrane features and proteins are known to be involved in the intrinsic resistance of some organisms by creating a barrier to the agent or encoding an efflux mechanism to prevent the agent from reaching its target (Russell, 2001). The importance of this membrane barrier could be a reason why many of the commercial decontamination agents contain surfactants. This implies that the intrinsic properties of a bacterium may be more important than its antibiotic-resistance status. This is supported by our finding that bleach was less effective against the antibiotic-sensitive molecular cloning strain Top 10 at holding times less than 240 min. This strain was also more resistant to Chemgene HLD4L than ATCC 25922.

Another important consideration is the ability of some bacterial species and isolates to produce biofilms. These are typically characterised by microcolonies of bacterial cells encased in an extracellular polymeric substance (EPS) matrix (Flemming et al., 2016). While we did not explore this in our study, many groups have previously shown that bacteria growing in a biofilm are more resistant to a variety of decontaminating agents when compared to planktonically-grown cells (Bridier et al., 2011).

## Conclusions

In conclusion, when deciding if a commercial decontamination agent is suitable for use, it would be sensible to consider how the standards used to test its efficacy relate to the application that the decontaminant is being considered for. As the responsibility for correct usage is with the end user, informed decisions regarding the choice of decontamination agent can only be made if those details are readily available, such as on the concentrate bottle. However, these details are often not available, or the recommendations on the label are different to how the product was tested. It is therefore imperative that manufacturers make the conditions of the testing available to product users. Considering our findings that species within the same genus and strains within a species can differ in their susceptibility to a variety of decontamination agents, we advise that users verify the efficacy of decontamination agents under the conditions in which they will be used to ensure that laboratory materials will be properly decontaminated. Our result show that merely increasing the recommended hold time rather than carrying out a proper validation is not sufficient for all organism-decontamination agent combinations.

## Acknowledgements

The authors would like to thank David Jenkins for his encouragement and support to carry out this study.

## References

Aronson, J.D. (1926). Spontaneous tuberculosis in salt water fish. The Journal of Infectious Diseases, 39(4), 315–320. https://doi.org/10.1093/infdis/39.4.315

Adair, F. W., Geftic, S. G., & Gelzer, J. (1969). Resistance of Pseudomonas to quaternary ammonium compounds. I. Growth in benzalkonium chloride solution. Applied microbiology, 18(3), 299–302. https://doi.org/10.1128/am.18.3.299-302.1969

Aires, J. R., Köhler, T., Nikaido, H., & Plésiat, P. (1999). Involvement of an active efflux system in the natural resistance of Pseudomonas aeruginosa to aminoglycosides. Antimicrobial agents and chemotherapy, 43(11), 2624–2628. https://doi.org/10.1128/AAC.43.11.2624

Bell, J. A., Brockmann, S. L., Feil, P., & Sackuvich, D. A. (1989). The effectiveness of two disinfectants on denture base acrylic resin with an organic load. The Journal of prosthetic dentistry, 61(5), 580–583. https://doi.org/10.1016/0022-3913(89)90280-1

Best, M., Sattar, S. A., Springthorpe, V. S., & Kennedy, M. E. (1990). Efficacies of selected disinfectants against Mycobacterium tuberculosis. Journal of clinical microbiology, 28(10), 2234–2239. https://doi.org/10.1128/jcm.28.10.2234-2239.1990

Cao, H., Lai, Y., Bougouffa, S., Xu, Z., & Yan, A. (2017). Comparative genome and transcriptome analysis reveals distinctive surface characteristics and unique physiological potentials of Pseudomonas aeruginosa ATCC 27853. BMC genomics, 18(1), 459. https://doi.org/10.1186/s12864-017-3842-z

Chapman JS (2003). Disinfectant resistance mechanisms, cross-resistance, and co-resistance. International Biodeterioration & Biodegradation, 51(4), 271–6.

Cuzon, G., Naas, T., & Nordmann, P. (2011). Functional characterization of Tn4401, a Tn3-based transposon involved in blaKPC gene mobilization. Antimicrobial agents and chemotherapy, 55(11), 5370–5373. https://doi.org/10.1128/AAC.05202-11

Freeman, J., Morris, A., Blackmore, T., Hammer, D., Munroe, S., & McKnight, L. (2007). Incidence of nontuberculous mycobacterial disease in New Zealand, 2004. The New Zealand medical journal, 120(1256), U2580.

Geraghty, R. J., Capes-Davis, A., Davis, J. M., Downward, J., Freshney, R. I., Knezevic, I., Lovell-Badge, R., Masters, J. R., Meredith, J., Stacey, G. N., Thraves, P., Vias, M., & Cancer Research UK (2014). Guidelines for the use of cell lines in biomedical research. British journal of cancer, 111(6), 1021–1046. https://doi.org/10.1038/bjc.2014.166

Griffiths, P. A., Babb, J. R., & Fraise, A. P. (1998). Mycobacterium terrae: a potential surrogate for Mycobacterium tuberculosis in a standard disinfectant test. The Journal of hospital infection, 38(3), 183–192. https://doi.org/10.1016/s0195-6701(98)90273-0

Korotetskiy, I. S., Joubert, M., Magabotha, S. M., Jumagaziyeva, A. B., Shilov, S. V., Suldina, N. A., Kenesheva, S. T., Yssel, A., Reva, O. N., & Ilin, A. I. (2020). Complete genome sequence of collection strain Acinetobacter baumannii ATCC BAA-1790, used as a model to study the antibiotic resistance reversion induced by iodine-containing complexes. Microbiology resource announcements, 9(3), e01467–19. https://doi.org/10.1128/MRA.01467-19

Makarova, O., Johnston, P., Walther, B., Rolff, J., & Roesler, U. (2017). Complete genome sequence of the disinfectant susceptibility testing reference strain Staphylococcus aureus subsp. aureus ATCC 6538. Genome announcements, 5(19), e00293–17. https://doi.org/10.1128/genomeA.00293-17

Minogue, T. D., Daligault, H. A., Davenport, K. W., Bishop-Lilly, K. A., Broomall, S. M., Bruce, D. C., Chain, P. S., Chertkov, O., Coyne, S. R., Freitas, T., Frey, K. G., Gibbons, H. S., Jaissle, J., Redden, C. L., Rosenzweig, C. N., Xu, Y., & Johnson, S. L. (2014). Complete genome assembly of Escherichia coli ATCC 25922, a serotype O6 reference strain. Genome announcements, 2(5), e00969–14. https://doi.org/10.1128/genomeA.00969-14

Oatway, W.H., & Steenken, W. (1936). The pathogenesis and fate of tubercle produced by dissociated variants of Tubercle Bacilli. The Journal of Infectious Diseases, 59, 306–325.

Russell A. D. (2001). Mechanisms of bacterial insusceptibility to biocides. American journal of infection control, 29(4), 259–261. https://doi.org/10.1067/mic.2001.115671

Russell, A. D., & McDonnell, G. (2000). Concentration: a major factor in studying biocidal action. The Journal of hospital infection, 44(1), 1–3. https://doi.org/10.1053/jhin.1999.0654

Rutala, W. A., & Weber, D. J. (1999). Infection control: the role of disinfection and sterilization. The Journal of hospital infection, 43 Suppl, S43–S55. https://doi.org/10.1016/s0195-6701(99)90065-8

Rutala, W.A., & Weber, D.J. (2015). Disinfection, Sterilization, and Control of Hospital Waste. In: Bennett, J.E., Dolin, r., & Blaser, M.J., ed. Mandell, Douglas, and Bennett’s Principles and Practice of Infectious Diseases. Eighth edition. Philadelphia: Saunders Elsevier, Volume 2, 3294–3309.

Snapper, S. B., Melton, R. E., Mustafa, S., Kieser, T., & Jacobs, W. R., Jr (1990). Isolation and characterization of efficient plasmid transformation mutants of Mycobacterium smegmatis. Molecular microbiology, 4(11), 1911–1919. https://doi.org/10.1111/j.1365-2958.1990.tb02040.x

Tattawasart, U., Maillard, J. Y., Furr, J. R., & Russell, A. D. (1999). Development of resistance to chlorhexidine diacetate and cetylpyridinium chloride in Pseudomonas stutzeri and changes in antibiotic susceptibility. The Journal of hospital infection, 42(3), 219–229. https://doi.org/10.1053/jhin.1999.0591

Trepanier, P., Mallard, K., Meunier, D., Pike, R., Brown, D., Ashby, J. P., Donaldson, H., Awad-El-Kariem, F. M., Balakrishnan, I., Cubbon, M., Chadwick, P. R., Doughton, M., Doughton, R., Hardiman, F., Harvey, G., Horner, C., Lee, J., Lewis, J., Loughrey, A., Manuel, R., … Woodford, N. (2017). Carbapenemase-producing Enterobacteriaceae in the UK: a national study (EuSCAPE-UK) on prevalence, incidence, laboratory detection methods and infection control measures. The Journal of antimicrobial chemotherapy, 72(2), 596–603. https://doi.org/10.1093/jac/dkw414

Wesgate, R., Grasha, P., & Maillard, J. Y. (2016). Use of a predictive protocol to measure the antimicrobial resistance risks associated with biocidal product usage. American journal of infection control, 44(4), 458–464. https://doi.org/10.1016/j.ajic.2015.11.009

